# Biocompatible polymeric microparticles serve as novel and reliable vehicles for exogenous hormone manipulations in passerines

**DOI:** 10.1101/2022.06.01.494300

**Authors:** Katharina Mahr, Maria Anzengruber, Anna Hellerschmid, Julia Slezaceck, Herbert Hoi, Guruprakash Subbiahdoss, Franz Gabor, Ádám Z. Lendvai

## Abstract

The administration of exogenous hormones emerged as an essential tool for field studies in endocrinology. However, working with wild animals remains challenging because under field conditions, not every available method meets the necessary requirements. Achieving a sustained elevation in hormone levels while simultaneously minimising handling time and invasiveness of the procedure is a difficult task in field endocrinology. Facing this challenge, we have investigated the suitability of biocompatible polymeric microparticles, a novel method for drug administration, as a tool to manipulate hormones in small songbirds. We chose the insulin-like growth factor -1 (IGF-1) as the target hormone because it receives great interest from the research community due to its important role in shaping life-history traits. Moreover, its short half-life and hydrophilic properties imply a major challenge in finding a suitable method to achieve a sustained, systemic long-term release. To study the release kinetics, we injected either IGF-1 loaded polylactic-co-glycolic acid (PLGA) microparticles or dispersion medium (control group) in the skin pocket of the interscapular region of captive bearded reedlings (*Panurus biarmicus*). We collected blood samples for 7 consecutive days plus an additional sampling period after two weeks and complemented these with an *in vitro* experiment. Our results show that *in vitro*, PLGA microparticles allowed a stable IGF-1 release for more than 15 days, following a burst release at the beginning of the measurement. *In vivo*, the initial burst was followed by a drop to still elevated levels in circulating IGF-1 until the effect vanished by 16 days post-treatment. This study is the first to describe PLGA-microparticles as a novel tool for exogenous hormone administration in a small passerine. We suggest that this method is highly suitable to achieve the systemic long-term release of hydrophilic hormones with a short half-life and reduces overall handling time, as it requires only one subcutaneous injection.

## Introduction

Exogenous hormone administration is a frequently used method to experimentally test for the ecological relevance of variations in hormone levels in different organisms (Crim 1985; Fusani 2008; Sopinka et al. 2015; Bonier & Cox 2020). Techniques that have previously been developed and used in laboratories (e.g., in pharmacology) recently became affordable and accessible to be safely applied to a variety of animal species in natural environments. The administration of hormones in a controlled setting (e.g., laboratory or housing facilities) allows standardised conditions (i.e., temperature, dietary regime, photoperiod), regular access to the target individuals, frequent handling, and strict monitoring of the treatment effects. On the contrary, in wild, free-ranging animals, experiments involving manipulations of hormone levels face several pitfalls and limitations (Fusani et al. 2005; Fusani 2008; Quispe et al. 2015; Ovid, Hayes & Bentley 2018).

There are different techniques for the administration of hormones available (e.g., oral, topical administration, intramuscular, subcutaneous or intraperitoneal injections/implants), but many cause only a transient elevation in hormone levels and/or require surgery (i.e., implantation) (Fusani 2008; Müller et al. 2009; Quispe et al. 2015; Ovid, Hayes & Bentley 2018; Vitousek et al. 2018). Many commonly applied hormones, including steroid hormones such as corticosterone and testosterone, have a short half-life, and sustained elevation requires daily, controlled oral or dermal application or injections (Sopinka et al. 2015). These methods are not feasible in the majority of field studies because repeated capture of target individuals is not possible or would seriously interfere with the natural behaviour of the animals (e.g., induce desertion, emigration from the site, interruption of mating or feeding behaviour) (Colwell et al. 1988; Putman 1995; Criscuolo 2001; Fusani et al. 2005; Ovid, Hayes & Bentley 2018). The work with wildlife underlies strict animal welfare regulations (Putman 1995; Gaunt et al. 1997; Fusani et al. 2005). Capture, restrain, sampling, the application of anesthesia, and the hormone manipulation itself might cause distress and have negative consequences on individuals living in natural conditions and consequently lead to a potential decrease in fitness and/or survival (Putman 1995; Fusani et al. 2005; Goutte et al. 2010; Spée et al. 2011; Bonier & Cox 2020; Huber et al. 2021). In addition, the impact of the application process itself and the effects of the hormone treatment (e.g., induced physiological responses, behavioural changes) might depend on the condition and life-history stage of an individual, both factors that are difficult to control for in field conditions (Colwell et al. 1988; Putman 1995; Criscuolo 2001; Ovid, Hayes & Bentley 2018). Finally, and contrary to many experiments in a clinical or preclinical setting, studies focusing on an ecological context of a given physiological response often aim to mimic hormone levels that correspond to a specific physiological phenotype within a population. The steady release of the hormone into the bloodstream and/or tissue is desirable to achieve hormone levels within the population’s estimated natural range (Fusani 2008; Quispe et al. 2015; Beck et al. 2016; Ovid, Hayes & Bentley 2018). Birds (in particular passerine species) have emerged as a popular study system because they display a large diversity of life-history traits, can be found all over the world as well in every habitat, and appear quite robust despite their often small size (Gaunt et al. 1997; Voss, Shutler & Werner 2010; Owen 2011). Hence, as Fusani (2008) reviewed, many pioneering studies on field endocrinology have been conducted on passerine species, and there are now a variety of safe options available to steadily release exogenous hormones over a specified timeframe (Beck et al. 2016). They have been tested sufficiently on passerine species in the field and the laboratory (Fusani 2008; Müller et al. 2009; Quispe et al. 2015; Ovid, Hayes & Bentley 2018; Vitousek et al. 2018).

One common method is the intraperitoneal or subcutaneous implantation of sterile, hormone-packed silastic tubes to facilitate the long-term release of a given hormone. They are reliable vehicles for hormone manipulations and cause long-lasting changes in peripheral hormone levels over several weeks. There are three frequently discussed downsides that include: i) an initial spike after implantation causing an elevation of the target hormone, sometimes far above natural levels, ii) silastic tubes are recommended to be removed after some time and therefore require repeated capture and handling, and iii) there are potential implications for animal welfare (i.e., capture stress or repeated wounding from surgery) (Horton, Long & Holberton 2007; Fusani 2008; Müller et al. 2009; Quispe et al. 2015). An alternative is the use of biodegradable time-release pellets designed to provide a steady release of hormones. However, these pellets were designed for small mammals, and their use in avian studies showed that their release kinetics in birds are characterized by a pronounced initial peak and a rapid depletion of the exogenous hormone, possibly due to higher avian body temperature and metabolism (Müller et al. 2009; Vágási et al. 2018). Another drawback is that they are only commercially available and are relatively expensive, but more cost-efficient/affordable alternatives have been recently developed using in-house made pellets from beeswax or hardened peanut oil (Quispe et al. 2015; Beck et al. 2016). In addition, and just like the silastic tubes, their chemical properties are only suitable for incorporating lipophilic hormones, such as steroids (e.g., Corticosterone, Testosterone). Yet many important peptide hormones are water-soluble and cannot be easily incorporated into the lipid matrix of the beeswax pellets. Another alternative technique that can achieve a sustained release of hormones are osmotic mini-pumps. These mini-pumps need to be implanted into the animal and function based on an osmotic pressure difference between a special osmotic layer within the pump and the surrounding tissue (Sinha & Trehan 2003). While these devices may control the release of theoretically any substance, due to their relatively large size and expensive price, they did not become widespread in field endocrinology (Pedersen & Saether 1999). An additional limitation of all pellets, tubes, and mini-pumps discussed above is that their initial implantation requires surgery (mini-pumps and silastic tubes even require additional surgery for removal), and the relatively rigid material and shape might constitute an irritation for the animal.

Here we test a novel, minimally invasive technique for exogenous hormone administration. We use a single subcutaneous injection of polylactic-co-glycolic acid (PLGA) microparticles to achieve a steady hormone release over several days. PLGA is a versatile co-polymer of lactic and glycolic acid that is clinically used as resorbable suture material and technologically for preparing drug delivery systems such as nano- or microparticles (Wischke & Schwendeman 2008). The use of PLGA for the protection and controlled delivery of proteins is well-established (Cohen et al. 1991; Kim, Chung & Park 2006; Dördelmann et al. 2014). It is a biodegradable and biocompatible polyester that degrades to non-toxic lactic acid and glycolic acid to finally yield carbon dioxide and water (Makadia & Siegel 2011). The degradation profile can be modified by the molecular weight of the polymer and the ratio of glycolic and lactic acid in the molecule (Park 1995). By entrapping drugs within a PLGA matrix, a continuous and controlled release profile can be achieved. Ideally, this reduces the frequency of drug administration and ensures a constant plasma level (Stevanović & Uskoković 2009). Furthermore, a biodegradable matrix material such as PLGA avoids the need for surgical removal of depleted delivery systems (Sinha & Trehan 2003).

We used insulin-like growth factor 1 (IGF-1) as the target substance. It is of increasing interest to many fields of the life sciences because it is an evolutionarily highly conserved metabolic hormone. It regulates a large array of physiological processes and has been shown to be associated with the development of major life-history traits (Dantzer & Swanson 2012; Lodjak & Verhulst 2020; Al-Samerria & Radovick 2021). One crucial aspect in working with IGF-1 is its short half-life in the circulation (in chicken, 32 minutes, regardless of the dose), which requires a suitable delivery system/vehicle to achieve a steady release and hence elevation of IGF-1 *in vivo* (Hameed et al. 2019; Sun, Yau & Chen 2019; Zhang et al. 2020). While a growing number of preclinical and clinical studies addressed this issue by using PLGA microparticles to increase systemic or local (i.e., specific target tissue) IGF-1 levels in laboratory- or farm animals (Lam et al. 2000; Yuksel et al. 2000; Meinel et al. 2001; Haney et al. 2018; Hameed et al. 2019; Zhang et al. 2020), this method has never been validated in a wild animal, let alone in a passerine species. To the best of our knowledge, only one experiment brought strong indication for the suitability of PLGA microparticles as a novel and minimally invasive tool to increase IGF-1 in the bearded reedling (*Panurus biarmicus*), a common, small (∼ 15 g) Eurasian songbird (Lendvai et al. 2021).

There is only a handful of studies examining the role of IGF-1 in regulating life-history traits of wild animal species, and even fewer testing the effects of an experimental increase of systemic IGF-1 on their physiology and behaviour (Dantzer & Swanson 2012; Lodjak & Verhulst 2020; Montoya et al. 2022). In the bearded reedling, IGF-1 has been studied in various contexts and is associated with regulating the stress response (Tóth, Ouyang & Lendvai 2018), ornament expression (Mahr et al. 2020) and feather moult (Lendvai et al. 2021). Lendvai et al. (2021) has shown that one single subcutaneous injection of PLGA microparticles loaded with IGF-1 resulted in a significant elevation of IGF-1 on day 1 post-injection and was sufficient to alter moult patterns and feather quality.

We now aim to investigate the release kinetics of PLGA microparticles loaded with IGF-1 in more detail, using captive bearded reedlings as models. The *in vivo* data will further be complemented with an *in vitro* study. The information we gain through this approach might provide novel and important insights into a promising method that increases hormone levels over the long term, while simultaneously avoiding supra-physiological hormone concentrations and decreasing handling time.

## Methods

### General methods and housing conditions

All birds were juveniles and originated from a large population of wild, free-living bearded reedlings (3 study sites in the Lake Neusiedl region, Burgenland, Austria: Winden am See 47°55’57.2”N 16°46’26.4”E; Jois 47°56’18.9”N 16°47’40.0”E; Breitenbrunn 47°55’25.4”N 16°45’29.9”E). They were caught using mist-nets and playback equipment. Immediately after the capture, a small blood sample was drawn. Since handling stress might affect circulating IGF-1 levels, the exact capture time (i.e., the bird hitting the net) was noted, and blood samples were drawn within 3 min. Subsequently, the birds were transferred to cloth cages provided with water and food (mealworms) and transported to the outdoor housing facilities of the KLIVV (Konrad Lorenz Institute of Ethology, Vienna, Austria).

They were housed in large outdoor aviaries, furnished with reed-bundles mimicking their natural environment. Food (fresh mealworms, insectivorous food containing quark, apples, carrots, dried insects, egg-food) and water were provided *ad libitum*. Moreover, the birds received a piece of apple, millet and vitamin supplements (Korvimin® ZVT + Reptil from WDT eG, Garbsen) weekly. Three days before the start of the experiment, birds were transferred into cages located in indoor housing facilities to allow a short acclimation period.

Each experimental cage contained 5 birds randomly assigned to the cages and experimental rooms. In this species separate housing is not recommended because reedlings are social animals and separation may lead to stress (Griggio & Hoi 2011; Tóth, Ouyang & Lendvai 2018). Although the juveniles were not displaying aggressive interactions, each cage was provided with 3 feeders and 2 water dispensers to guarantee sufficient access to food and water. The indoor facilities were equipped with a ventilation system that allowed natural airflow, with temperatures resembling ambient climatic conditions. Similarly, light conditions were adjusted to the current conditions in the natural environment (light between 0600 and 1930).

### IGF-1 Microspheres preparation

Poly-(lactide-co-glycolide) RG502H (PLGA) was obtained from Evonik Nutrition & Care GmbH (Essen, Germany). Human recombinant insulin-like growth factor 1 (IGF-1) was bought from PreproTech EC, Ltd. (London, UK). Polyvinylalcohol (PVA, 87-90% hydrolysed, 30-70 kDa), dichloromethane (DCM) and all further chemicals were purchased from Carl Roth GmbH (Karlsruhe, Germany) or Sigma Aldrich (St. Louis, MO, USA).

Microparticles were prepared by a w1/o/w2 double emulsion technique with modifications as initially described by Meinel et al. (2001). The inner aqueous phase (w1) consisted of 20 µg IGF-1, 10 mM sodium-succinate and 140 mM NaCl dissolved in 150 µl purified water. Additionally, 2.5 mg bovine serum albumin and 2.0 mg succinylated gelatine (Gelofusin®) were added to stabilize IGF-1 during particle preparation and to control release. The efficacy of those compounds was previously investigated by Meinel et al. (2001). The aqueous phase w1 was then injected into a solution of 50 mg PLGA RG502H in 2 ml DCM and emulsified via ultrasonication using a Bandelin HD70 Sonoplus sonifier for 15 seconds at 30% amplitude to yield emulsion 1. Subsequently, emulsion 1 was added to 30 ml 5% aqueous PVA solution (w/v) and stirred for one minute at 500 rpm yielding the final w1/o/w2 double emulsion. For particle hardening, the final dispersion was diluted with 400 ml of purified water and stirred for 30 minutes at 100 rpm and under constant airflow to ensure complete evaporation of DCM. The microparticles were washed three times with purified water and centrifugation at 1068 × g for 10 minutes. The purified microparticles were freeze-dried for at least 24 hours and stored at 4°C until further use.

Prior to animal application, IGF-1 loaded microparticles were suspended in a dispersion medium yielding a final IGF-1 concentration of 280 ng/100 µl. The dispersion medium consisted of 1.5% (w/w) carboxymethylcellulose, 5% (w/w) mannitol and 0.02% (w/w) polysorbate 80 dissolved in a sterile, physiologic sodium chloride solution.

### Characterization of IGF-1 containing PLGA-microparticles and Quantification of IGF-1 by HPLC analysis

The particle size distribution of the microparticle suspension was determined by laser diffraction using a Mastersizer 3000 with a Hydro SV dispersion unit (Malvern Instruments, Malvern, UK). Lyophilised microparticles were suspended in purified water prior to particle size characterisation. The suspension was stirred at 700 rpm during the measurement. Further, particle size and morphology were analysed via light microscopy using a Zeiss Epifluorescence Axio Observer.Z1 deconvolution microscopy system.

Scanning electron microscopy was performed to examine the morphology of PLGA microparticles. The lyophilised microparticles were placed on aluminium SEM stub and high vacuum secondary electron imaging was performed using an Apreo VS SEM (Thermo Scientific, The Netherlands) at 1.0 kV.

IGF-1 was quantified using a Shimadzu Nexera XR HPLC (Shimadzu, Kyoto, Japan) equipped with an analytical CN-RP column (NUCLEODUR 100-5, CN-RP, 150 × 4.6, 5 µm column; Macherey-Nagel GmbH & Co. KG, Düren, Germany) and a UV-diode-array detector. A gradient elution protocol with acetonitrile (ACN) and purified water at a 0.8 ml/min flow rate was employed. Formic acid was added to both solvents at a concentration of 0.1%. Over ten minutes, the eluent was changed from 5% ACN and 95% water to 45% ACN and 55% water. IGF-1 was detected at 214 nm and the amount was calculated from a standard curve. To quantify encapsulated IGF-1, a certain amount of particles was dissolved in 1 ml of DCM + acetone (3+7). The sample was then centrifuged at 20817 × g for five minutes and the supernatant was discarded. This purification procedure was repeated three times. The final residue was dissolved in 0.1 M acetic acid and analysed by HPLC as described above.

For in-vitro IGF-1 release experiments, a dispersion of 40.0 mg microparticles in one ml PBS with Ca^2+^/Mg^2+^ pH 7.4 was incubated under constant agitation at 37°C. The IGF-1 concentration in the acceptor medium was determined in regular intervals. At each time point, 200 µl of the acceptor medium were withdrawn and replaced by fresh PBS buffer. The aliquot was lyophilized, dissolved in 0.1 M acetic acid, and subjected to HPLC analysis.

### Experimental procedure

After an acclimation period of 48 hours, a small blood sample was collected from all individuals between 0900-1200. Similarly to the sampling in field conditions, the exact time between entering the experimental room (4 rooms with separate entrances) and drawing a blood sample was noted to correct for a potential effect of sampling/handling time on circulating IGF-1 levels. Samples were immediately stored in a fridge until further processing. Subsequently, the IGF-1 treatment (IGF-1) or the control (C, dispersion medium without IGF-1) were applied. Birds were randomly assigned to treatments and each cage contained individuals from both treatment groups. To apply the exogenous hormone manipulation, a spot in the interscapular region was cleared with alcohol swabs (Rosner GmBh, Vienna) and 100 μl of a dispersion containing either microsphere loaded with recombinant human IGF-1 or only the dispersion medium were injected. For the application, syringes without residual volume (B Braun™ Omnifix®-F Spritzen) and disposable sterile needles (21 G, B Braun™) were used. The applied liquid is highly viscous and forms a depot from where the active ingredient is continuously released.

In order to monitor the hormonal changes after the manipulation, blood samples were collected from different birds on different days over a period of 7 days. Thereby birds were grouped into 6 cohorts. Each cohort consisted of 4 birds from the control group and 6 birds from the treatment (IGF-1 injection) group. Starting with the first cohort, a blood sample was collected from a different cohort every 24 hours (± 1 h) until day 6; on day 7 and 16 post-injection a subset of birds was sampled again.

After each sampling unit, the injection site was examined carefully to record any potential inflammation or skin reaction. The liquid distributed evenly under the skin and the birds’ movements were not restricted. The skin around the injection site did not show any signs of inflammation or necrotic tissue.

### IGF1-1 Assays

Plasma IGF-1 levels were measured by a competitive ELISA at the University of Debrecen described previously in Mahr et al. (2020). Briefly, 96-well microplates were coated at 4°C overnight with 100 µl of an antibody raised against IGF-1 in rabbits. The coated plate was incubated for 2 h at room temperature with either 20 µl of standard (known concentrations of synthetic chicken IGF-1 in serial dilutions starting at 500 ng/ml) or 20 µl of sample and 100 µl of biotinylated IGF-1 as a tracer. After incubation, the microplate was washed three times with 250 µl of PBS containing 0.025% Tween 20. After washing, 100 µl of streptavidin-horseradish peroxidase conjugate was added to all wells and incubated at room temperature for 30 min. After washing, 100 µl of tetra-methyl-benzidine was added to the wells and incubated for 30 min. The enzymatic reaction was stopped by adding 100 µl of 1M H_2_SO_4_, and optical density was measured at 450 nm (reference at 620 nm) using a Tecan F50 microplate reader.

### Statistical analyses

In total samples from 60 individuals were collected and analysed (IGF-1, n = 37; Control, n = 23). Results from *in vitro* release experiments were analysed using GraphPad Prism 9. All data from *in vitro* release experiments are presented as mean with n ≥3 and ± standard deviation. The release kinetics was determined by fitting the drug release data to the pharmacokinetic models of the first order, Higuchi, Korsmeyer-Peppas and Weibull model (Costa & Sousa Lobo 2001; Siepmann & Siepmann 2008). The adjusted coefficient of determination (R^2^ adjusted) was calculated to evaluate the fit of the release data to the respective pharmacokinetic model (Costa & Sousa Lobo 2001).

We used general linear mixed models implemented in the packages lmerTest in the R version ‘Bird Hippie’ (4.1.2) (R Core Team 2021) to analyse variation in the *in vivo* IGF-1 plasma concentrations, with time as a fixed factor and random intercepts for individual birds to control for the repeated measurements. Within each sampling unit, treatment groups were compared using post-hoc contrasts, as implemented in the package emmeans (Lenth 2022).

## Results

### Characterization of microparticles and in vitro release of IGF-1

Microparticles exhibited a spherical shape with a structured surface and a mean volumetric particle diameter D[4;3] of 28.4 µm ± 0.39, a Dx(90) of 51.3 µm ± 0.85 and a Dx(10) of 10.4 µm ± 0.12. These results were corroborated by the images of the microscopic analysis. An additional SEM analysis of the particles revealed a highly porous particle structure created due to the porogens sodium chloride and sodium succinate during the particle preparation process (Qutachi et al. 2014) (Fig. 1). The encapsulation efficiency of IGF-1 into the microparticle matrix amounted to 70-80% and was calculated in reference to the maximum loading capacity of 20 µg IGF-1 per 50 mg microparticles.

**Figure 1.**
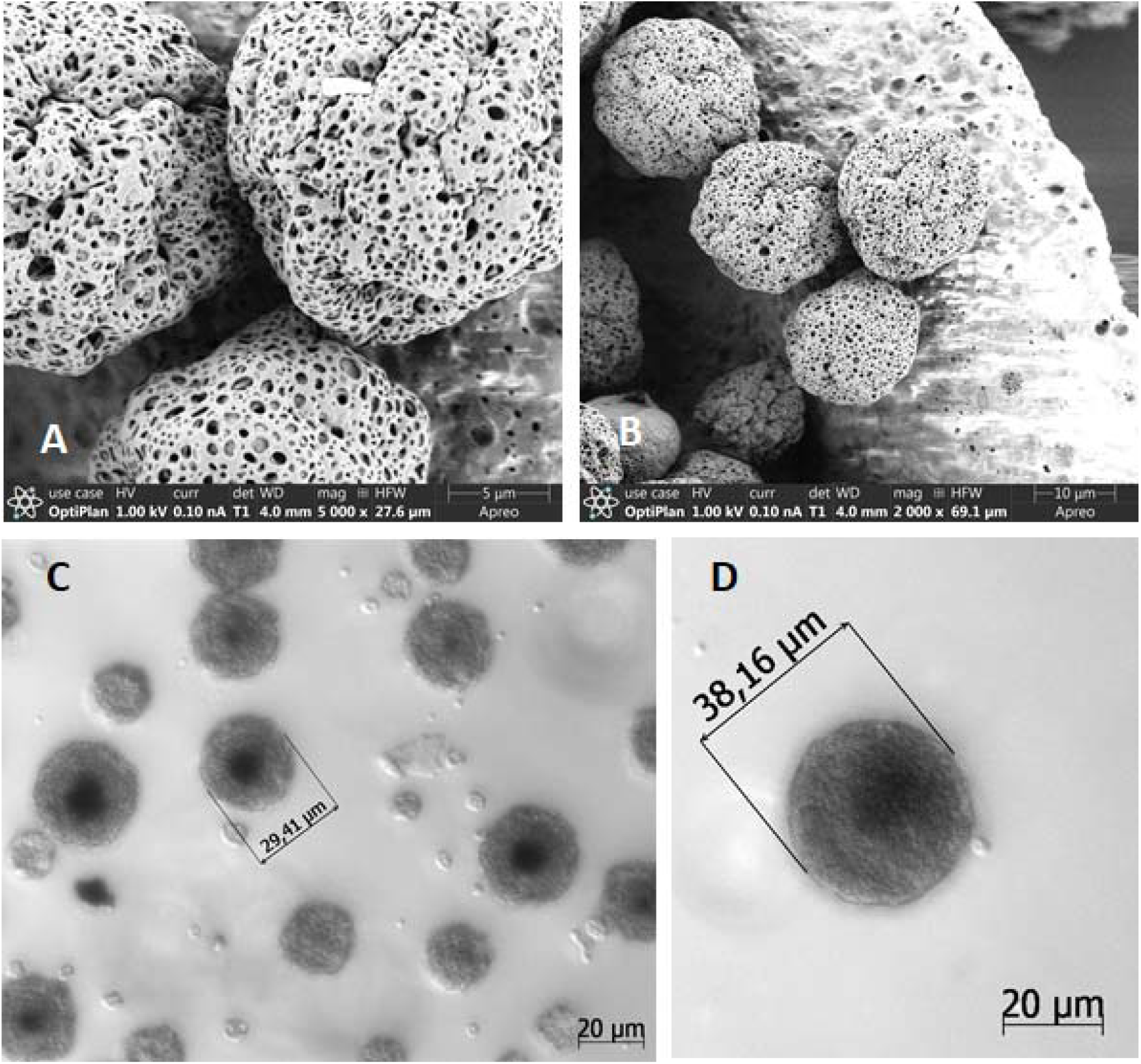
Scanning electron microscopy images (A and B) and light microscopic images of PLGA RG502H microparticles containing IGF-1 (C and D).

The release experiments revealed an initial burst release within the first 24 hours, amounting to around 40% of the payload, followed by a continuous release of IGF-1 from the microspheres over the next 14 days. After 15 days, no further significant release of IGF-1 was observed (Fig. 2). These results are in accordance with the release patterns previously presented by Meinel et al. (2001). The release kinetics of IGF-1 from PLGA microparticles correlated best with the pharmacokinetic model of Korsmeyer-Peppas and that of Weibull, with an R^2^ adjusted of 0.958 and 0.934, respectively. The drug release kinetic that follows the burst release fitted well to first-order kinetics as well (R^2^ adjusted 0.943), underpinning the gradual, almost linear release of IGF-1 from the particle matrix after the initial burst.

**Figure 2.**
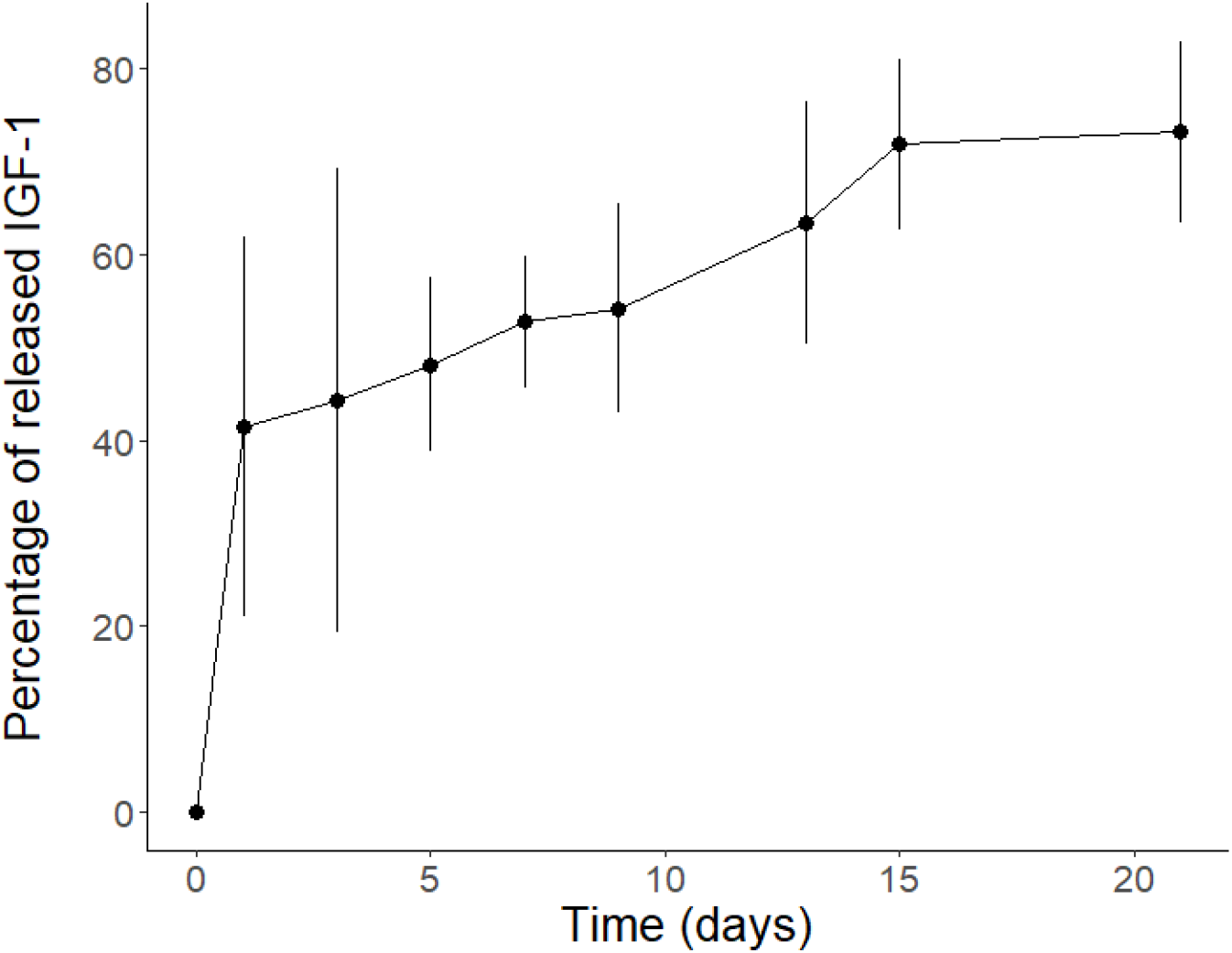
Cumulative release (mean ± SD) *in vitro* of IGF-1 from PLGA RG502H microparticles over time.

### In vivo release of IGF-1

Before the treatment, there was no difference in IGF-1 levels between the experimental groups (Field: t = 0.65, p = 0.51, day 0: t = 0.63, p = 0.52). According to our expectations, IGF-1 levels increased markedly the day following the injections in the treatment group, but surprisingly, there was also an initial increase in IGF-1 levels in the control group. However, the increase in the treatment group was stronger, resulting in a significant difference between the experimental groups on day 1 (t = 2.21, p = 0.02. Fig. 3). After this initial peak, IGF-1 concentrations dropped in both groups over the following days, and the difference between the treatment and control group remained significant or at the boundaries of statistical significance (day 2: t = 1.65, 0.099, day 3: t = 1.76, p = 0.079, day 5: t = 2.03, p =0.04, day 7: t = 2.50, p= 0.01). Interestingly, on day 4 and 6 the experimental groups converged statistically (day 4: t = 1.13, p = 0.26, day 6: t = 0.64, p = 0.51). Finally, by day 16, IGF-1 levels in the treatment group fell back to the levels of the controls (t = 0.44, p = 0.66).

**Figure 3.**
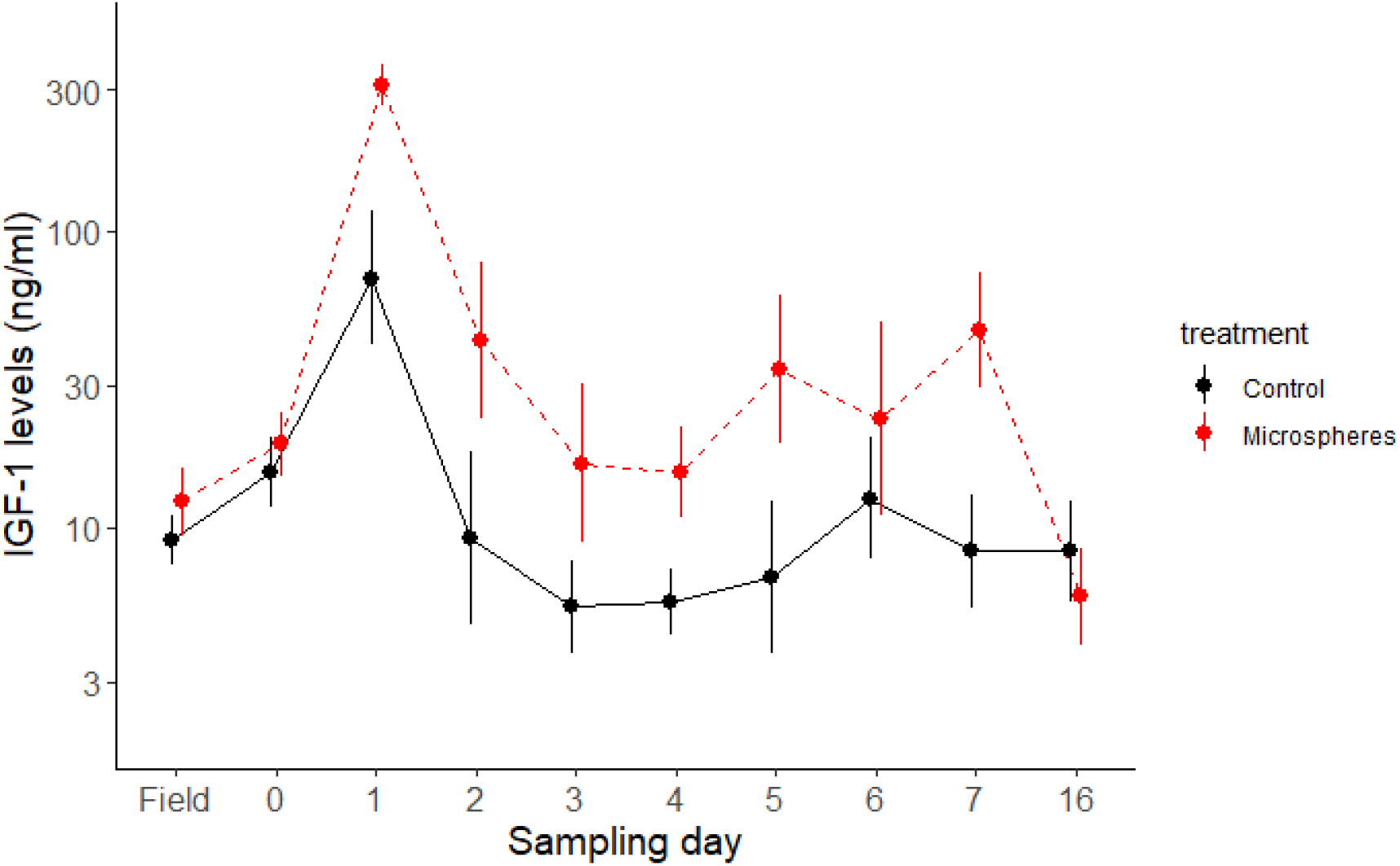
A single subcutaneous injection of IGF-1 loaded PLGA microparticles (day 0) resulted in a sustained elevation of circulating IGF-1 concentrations in bearded reedlings.

## Discussion

In this experiment, we tested the suitability of polymeric microparticles as biocompatible vehicles for the elevation of systemic IGF-1 in captive bearded reedlings over a period of 16 days. Our results show that a single subcutaneous injection of IGF-1-loaded PLGA microparticles resulted in an initial spike (i.e., burst release) of IGF-1 with hormone levels remaining significantly elevated for more than 7 - and less than 16 days. Overall, the *in vivo* release dynamics correlate well with the *in vitro* release profile and resemble previous experiments (Lam et al. 2000; Meinel et al. 2001). The *in vitro* release assay involves no degradative processes of the incorporated medical cargo; therefore a decrease in the IGF-1 concentration would indicate degradation or instability of IGF-1 in the release medium. A rising concentration *in vitro* indicates a release of IGF-1 from the particles. This can be observed until day 15, whereas between day 15 and day 20, there is no further change in the measured IGF-1 concentration *in vitro*; therefore, no additional IGF-1 is released. *In vivo*, the administered IGF-1, however, is degraded and used up over time. If degradative processes are faster than the continuous release from the particle matrix, IGF-1 concentration decrease *in vivo*. Accordingly, the IGF-1 concentration *in vivo* drops back to physiological levels as soon as no further IGF-1 is released from the particle matrix. A slightly prolonged drug release was yet observed in the *in vitro* setting compared to *in vivo*. This effect has been described previously and is due to faster degradation of the polymers *in vivo* (Cleland et al. 1997; Lam et al. 2000).

Both the *in vivo* and *in vitro* experiments are characterised by an initial burst release within the first 24 h followed by a continuous release over the next two weeks. This heterogeneous release profile hints towards the involvement of different mechanisms in drug release processes from PLGA microparticles (Xu et al. 2017). The disintegration of the particles and the formation of cracks on the PLGA matrix (i.e., bulk-erosion) and the subsequent dissolution of drug molecules near the particle surface explain the initial burst release. This fast drug release is then followed by a period of slower diffusion based, liberation of the encapsulated IGF-1 (Meinel et al. 2001; Makadia & Siegel 2011; Han et al. 2016). Although, in the present experiment, IGF-1 concentrations remained in the physiological range, as some control birds and individuals before the treatment had high IGF-1 values comparable with the day-1 peak of treated birds, the question arises how an initial burst release after injection can be avoided in future studies. This is an unwanted side effect of many exogenous hormone manipulations and may lead to an overshoot of the target hormone to levels far above the physiological range (Fusani 2008; Quispe et al. 2015). Previous experiments on rats (Lam et al. 2000) indicate that this effect can be avoided by initiating a “pre-burst” of PLGA microparticles in the injection vehicle. However, it should be considered that birds have higher body temperatures (∼41°C) than rats (37.5-38.5°C) (Gudjonsson 1932) and it has been shown that the release kinetics of the polymers are sensitive to temperature. Hence, this process might not be as effective for avian species and only lead to faster depletion of the encapsulated peptide.

In the *in vivo* experiment, we observed differences in IGF-1 levels between the control and treatment birds up to day 7, albeit on days 4 and 6 post-injection, the difference between the treatment groups became non-significant. This result is consistent with Lendvai et al. (2021), where IGF-1 concentrations were significantly elevated 24h after the microsphere injection but became similar to the control group by day 4. While there may be a decrease in the hormone-releasing efficiency around day 4, this decrease may also be due to the initially larger variance of release kinetics at this stage, which also becomes visible in the *in vitro* study (Fig. 2). Our results also show that even if there is a genuine decrease of IGF-1 at this stage, it is temporary, as by day 5 and 7 the IGF-1 concentrations differed significantly between the control and treatment groups (Fig. 3). However, despite this pattern, a clear trend indicates that overall, IGF-1 levels remained higher in the experimental than in the control group (Fig. 3). While these differences are sometimes marginally non-significant, this is most likely due to low statistical power resulting from repeated sampling, the small number of individuals we sampled each day, and the statistical correction for a higher number of multiple comparisons. When we pooled two consecutive sampling days following day 1, the overall differences between IGF-1 and control birds remained statistically significant throughout the first week up to day 7 (day 2-3: p = 0.02, day 4-5: p = 0.03, day 6-7: p = 0.02).

An unexpected result showed a pronounced peak in IGF-1 levels on day 1 post-injection in both experimental groups (i.e., also control birds). While this effect resembled the findings of Lendvai et al. 2021, the observed elevation was more pronounced here. All birds were captured on day 0, and after an initial blood sample, they were injected with the respective treatment (control/IGF-1) and measured for various metrics (not included in this study). This resulted in multiple handling events and an overall longer procedure. On subsequent days, only a subset of birds was captured, and apart from a rapid blood sample, only body mass was recorded. Therefore, this initial spike at IGF-1 levels might be the carry-over effect of the arguably stronger disturbance (e.g., long handling time) on the previous day. IGF-1 levels have been shown to decrease due to stressful stimuli (Tóth, Ouyang & Lendvai 2018; Valenzuela et al. 2018; Vágási et al. 2020), but there is strong indication that IGF-1 levels increase over the long term to minimise detrimental effects of chronic stress (McCormick et al. 1998; Xin et al. 2018), which might explain the initial increase of IGF-1 in control birds. IGF-1 is also known to regulate immunity, wound healing and inflammatory processes (Bos et al. 2001; Semenova et al. 2008; Emmerson et al. 2012). Hence, an alternative explanation for the temporary increase of IGF-1 might be due to the injection itself. During the experiment, we controlled the injection site daily to record any adverse effects of the procedure. Even though no inflammation or necrotic tissue was detected, the injection of IGF-1 loaded PLGA microparticles or dispersion medium itself might cause a temporal irritation at the injection site. In conclusion, it is highly likely that both the effects of blood collection and handling on the physiological stress response and the local immunological processes that may have been triggered by the injection could be responsible for the observed pattern.

Some techniques for exogenous hormone manipulations (e.g., silastic tubes) allow a continuous release of a target hormone for extended time (up to 4 weeks) (Fusani et al. 2005; Ovid, Hayes & Bentley 2018), but in this study, we have only achieved an increase of systemic IGF-1 levels for a period of at least 7, but less than 16 days. However, non-biodegradable pellets or silicone tubes require surgery to replace the vehicle after depletion (Ovid, Hayes & Bentley 2018), and biocompatible polymeric microparticles require only an additional subcutaneous injection to achieve a prolonged release. When wild animals cannot be easily recaptured, adjusting the properties of the PLGA microparticles (e.g., size or resomer type) may offer a possibility to modify the release kinetics and achieve even longer-lasting hormone treatments (Sun et al. 2008; Busatto et al. 2018; Matejkova & Podhorec 2019; Patel, Jha & Patel 2021).

## Conclusion

The application of IGF-1 loaded PLGA microparticles requires only a few handling steps while allowing a continuous release of the target hormone for several days and potentially weeks. In addition, we did not observe any adverse effects, such as inflammation or necrotic tissue on the injection site or noted any losses of birds within the course of the study. Our results provide further confirmation for the versatility of PLGA microparticles. Considering their suitability for the application of a variety of substances (e.g., steroid hormones) with different chemical properties, this technique is a highly promising resource that will open new possibilities for field studies in physiology and behaviour.

## Acknowledgements

We would like to thank A. Hloch, J. Cornils, F. Hölzl, S. Graf, and A. Aispuro for their help and support during the data collection, and A. Haw and N. Huber for supervising the experimental procedure and providing veterinary care and advice. We are also grateful for the advice and initial trials from B. Gander and S. Kindgen-Vogel. A thank you also goes to the animal caretakers (N. Skupa and S. Mali) for their work and to R. Hengsberger for copy-editing the manuscript.

The study was financed by the Erwin Schro□dinger fellowship (J4235-B29) granted by the FWF (Austrian Science Fund). During writing ÁZL, HH and KM were additionally supported by the Austrian Agency for International Cooperation in Education and Research (OeAD; WTZ grant (HU 05/2020)). ÁZL was supported by grants from the National Development, Research and Innovation Office (2019-2.1.11-TÉT-2019-00043 and OTKA K139021)

All procedures were in accordance with the ethical standards of the institution and were approved by the Austrian Federal Ministry for Education, Science and Research (§26 des Tierversuchsgesetzes 2012-TG 2012: A4/NN.AB-10087-22-2020; GZ 2020-0.292.788 and GZ 2020-0.466.191.).

## Conflict of interest

There is no conflict of interest.

## Author contributions

KM, ÁZL and MA conceived and conducted the experiment, KM, AH, JS and MA collected the samples and the data, ÁZL and MA measured the samples, FG and ÁZL contributed reagents, ÁZL, MA and GS analysed the data, KM, ÁZL and MA wrote the article, with significant contribution from JS, FG, AH and HH. All authors contributed critically to the drafts and gave final approval for publication

## Data availability

Data will be uploaded upon request.

## Notes

### Competing Interest Statement

The authors have declared no competing interest.

